# Deletion of Small Ubiquitin-like Modifier-1 Attenuates Behavioral and Anatomical Deficits by Enhancing Functional Autophagic Activities in Huntington Disease

**DOI:** 10.1101/2021.02.15.431277

**Authors:** Uri Nimrod Ramírez-Jarquín, Manish Sharma, Neelam Shahani, Srinivasa Subramaniam

## Abstract

Mutant HTT (mHTT) associated with Huntington disease (HD) affects the central nervous system by prominent atrophy in the striatum and promotes psychiatric, cognitive, and choreiform movements, although the exact mechanism remains obscure. Previous studies have shown that SUMO1 (Small Ubiquitin-like Modifier-1) modification of mHTT promotes cellular toxicity, but the in vivo role and functions of SUMO1 in HD pathogenesis are unclear. Here, we report that SUMO1 deletion in Q175DN HD-het knock-in mice (HD mice) prevented age-dependent HD-like motor and neurological impairments and suppressed the striatal atrophy and inflammatory response. SUMO1 deletion caused a drastic reduction in soluble mHtt levels and nuclear and extracellular mHtt inclusions, while increasing cytoplasmic inclusions in the striatum of HD mice. SUMO1 deletion also enhanced autophagic activity, characterized by augmented interactions between mHTT inclusions and a lysosomal marker (LAMP1), increased LC3B/LAMP1 interaction, and decreased sequestosome-1 (p62) and mHTT and diminished p62/LAMP1 interactions in DARPP-32–positive medium spiny neurons (MSNs) in HD mice. Depletion of SUMO1 in an HD cell model also diminished the mHtt levels and enhanced autophagy flux. In addition, the SUMOylation inhibitor ginkgolic acid strongly enhanced autophagy and diminished mHTT levels in human HD fibroblasts. These results indicate that SUMO is a critical therapeutic target in HD and that blocking SUMO may ameliorate HD pathogenesis by improving autophagy activities.

## INTRODUCTION

Expansion of the CAG repeat in the huntingtin (HTT) gene causes the motor disturbance, cognitive loss, and psychiatric manifestations associated with Huntington Disease (HD). This mutant HTT (mHTT) affects multiple signaling and pathways, including vesicle- and microtubule-associated protein/organelle transport, transcription, and calcium, sphingosine, cysteine, and p53/GAPDH signaling, as well as functional connectivity, secretory pathway function, cell division, mitochondrial abnormalities, and ribosome stalling [1-23]. However, the exact mechanism by which mHTT induces its pathological effects in the brain remains unclear.

Multiple lines of evidence point to an aberration in autophagy—an essential degradative pathway that maintains neuronal homeostasis—as the root cause of HD [24-28]. Studies have implicated aberrant autophagy in HD [27, 28], as well as a failure of autophagic processes such as cargo-loading, in the accumulation of mHTT and the progression of the disease [24, 25, 29]. Normal HTT serves as an autophagy scaffold and interacts with the Atg8 homologs, ULK1 and p62, during autophagy [30, 31] but the molecular mechanism and the precise role of mHTT in autophagy dysregulation remain elusive [32]. Accordingly, molecular and pharmacological agents that can promote mTOR-dependent and independent-mechanisms of autophagy are being actively tested in preclinical models [33-55]. Nevertheless, the actual alterations in autophagy homeostasis and the treatments that can lead to its restoration are unknown [32, 56]. This gap in knowledge remains a major hurdle in identifying targets for successful treatment of HD.

One potential target is the small-ubiquitin-like modifier (SUMO), which consists of three ubiquitously expressed paralogs in vertebrates: SUMO1, SUMO2, and SUMO3. It is a conserved ∼10.5kDa protein modifier that attaches covalently to lysine residues on multiple substrate proteins in a dynamic and reversible manner [57] in a process known as SUMOylation. Protein modification by SUMO is implicated in a diverse array of neurological disorders [58-62] and is known to regulate a wide range of cellular processes, including gene expression, nuclear transport, signal transduction, apoptosis, and autophagy [63-77]. Experiments with null mice confirm that SUMO1 and SUMO3 are indispensable for development, while deletion of SUMO2 is embryonically lethal[78-80]. These studies indicate critical roles for SUMO in an organism’s growth and development, but its mechanistic connection to neurodegenerative disease is less clear.

We have reported that a striatal enriched protein, Rhes, is SUMO modified on multiple lysine residues and serves as a physiological regulator of SUMOylation and gene transcription [81, 82]. Rhes interacts with mHTT and modifies it by SUMO1, consistent with other reports [83, 84]. SUMO1 modification of mHTT increases the solubility of mHTT, enhances Rhes-mediated transport of vesicle-bound mHTT via tunneling-like nanotubes [85], and promotes cellular toxicity [86]. Accordingly, Rhes deletion ameliorates, and Rhes overexpression worsens, the HD phenotype in HD cell models (primary neurons, mouse cells, and hESC-derived medium spiny neurons [MSNs]) and HD mouse models [86-94]. Recently, Rhes has been linked to tau-mediated pathology involving SUMOylation, and Rhes alterations are now identified as a novel hallmark of tauopathies [95, 96]. These studies indicate that Rhes–SUMO signaling pathways participate in neurodegenerative disease processes by regulating disease-relevant proteins.

However, no mechanisms or systematic evaluations have been identified to explain how SUMO might determine the course of neuronal decline and eventual striatal atrophy in slowly progressing knock-in HD mouse models. Here, we deleted SUMO1 in the Q175DN HD-het (Q175DN) knock-in mouse model of HD [97], which contains the human HTT exon 1 with expanded CAG repeats, and we characterized the resulting Q175DN-SUMO1KO mouse line longitudinally to determine the role of SUMO1 in terms of the nature, extent, and age of onset of behavioral deficits. We also performed anatomical, molecular, and biochemical signaling of the striatal tissues. Finally, we evaluated the effect of SUMO1 deletion or pharmacological SUMO inhibition on biochemical signaling by using HD mouse and human cell culture models.

## RESULTS

### SUMO1 deletion diminishes HD-associated behavioral deficits in Q175DN mice

To determine the role of SUMO1 in HD pathogenesis using HD knock-in mice [neomycin-deleted Q175DN HD-het (Q175DN)] mice [97], we generated HD mice lacking a SUMO1 gene (Q175DN-SUMO1KO). We included mice of both genders at a mixed ratio in our longitudinal analysis (Fig. 1A, Table 1). A slight decrease in body weight was noted at about 10 months of age in the Q175DN mice (n = 28), consistent with previous reports[97-99], also in the Q175DN-SUMO1KO mice (n = 30) when compared to the control groups (WT, n = 37 and SUMO1KO, n = 27), indicating that SUMO1 does not modulate the body weight loss observed in HD (Fig. 1B). However, the Q175DN-SUMO1KO mice did not show the progressively worsening rotarod behavior deficits that were observed in the Q175DN mice beginning at 8-10 months of age [97] (Fig. 1C). The Q175DN mice also showed deficits in fine motor coordination and balance in beam walking, but these deficits were also diminished in the Q175DN-SUMO1KO mice (Fig. 1D). Open-field tests revealed age-dependent hypoactivity in the Q175DN mice compared to WT mice, whereas the Q175DN-SUMO1KO mice showed significantly less hypoactivity compared to Q175DN at 8 months but not in later months (Fig. 1E). We also employed a battery of behavioral tests to quantify the neurological dysfunction. These tests include walking on a ledge, clasping, gait, kyphosis (spine curvature), and tremor, which can be averaged as a ‘composite’ score [100]. Behavioral battery tests revealed that Q175DN mice performed worst on ledge, clasping, gait, kyphosis, and tremor testing, but the performance of these tasks was significantly better in Q175DN-SUMO1KO mice in both individual tests and composite scores (Fig. 1F, G). Thus, SUMO1 deletion attenuated the severity of behavioral and neurological abnormalities associated with HD.

**Table 1:** Details of mice

| Genotype | Male | Female | Total |
| --- | --- | --- | --- |
| WT | 22 | 15 | 37 |
| Q175DN | 19 | 9 | 28 |
| Q175DN-SUMO1KO | 13 | 17 | 30 |
| SUMO1KO | 15 | 12 | 27 |

**Figure 1.**
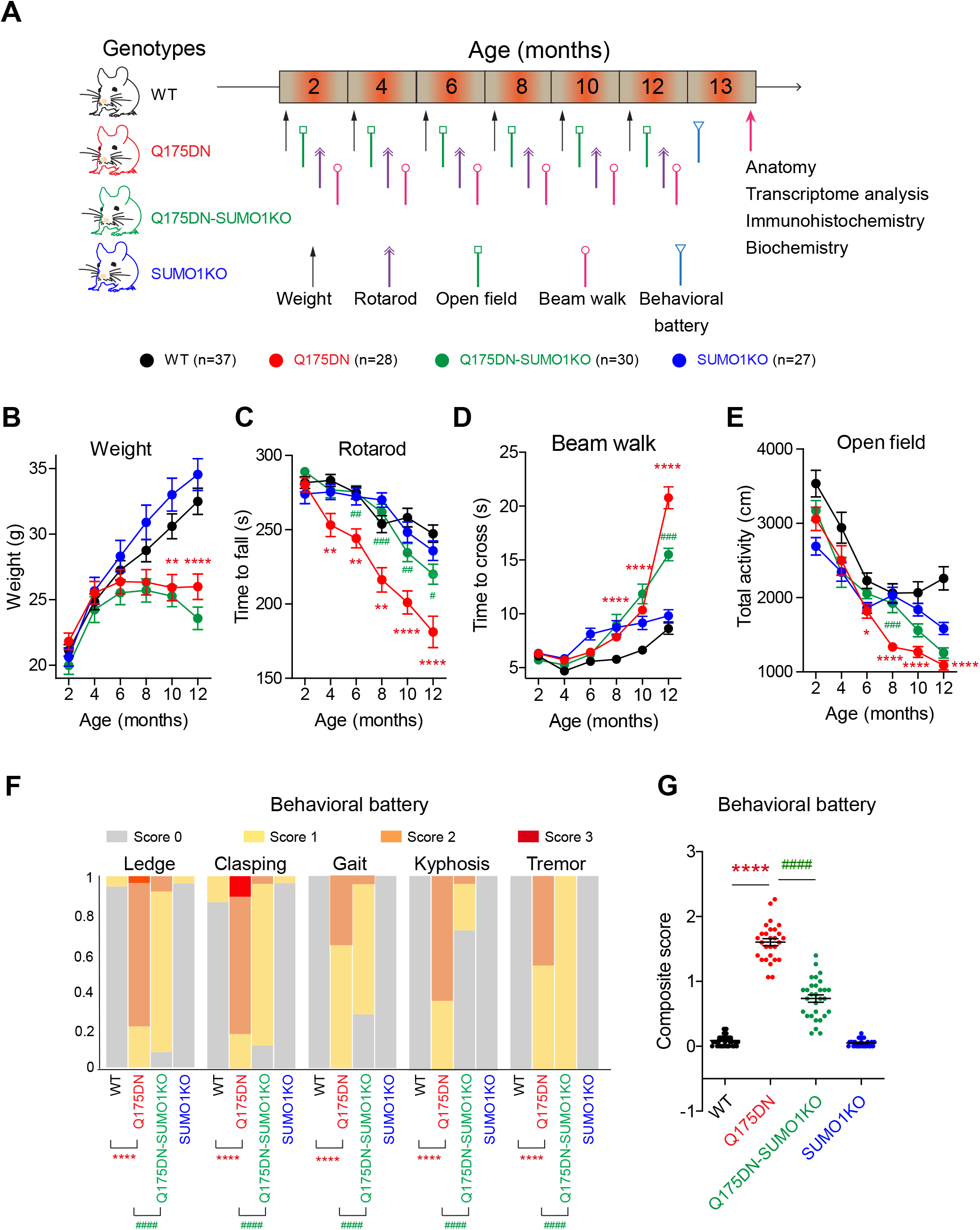
Effect of SUMO1 deletion on age-associated motor abnormalities in Q175DN knock-in HD mice model. (A) Shows the genotypes, timeline and experimental design for the animal study. (B-G) Body weight and behavioral analysis at 2-12 months of age; Body weight (B), rotarod (C), beam walk (D) and open field (E), behavioral battery (F), behavioral battery composite score (G). Behavioral batteries included ledge-test, clasping, gait, kyphosis, and tremor evaluation. The frequency of each behavior according to the genotype is shown in F, a composite score for all the behavioral battery is shown in G. Data are mean ± SEM (WT, n = 37, Q175DN, n = 28, Q175DN-SUMO1KO, n =30, SUMO1KO, n = 27). \**P* < 0.05, \*\*\*\**P* < 0.0001 between Q175DN and WT; ^#^*P* < 0.05, ^##^*P* < 0.01, and ^###^*P* < 0.001 between Q175DN-SUMO1KO and Q175DN by two-way ANOVA mixed-effects model (REML) followed by Tukey’s multiple comparison test (For B-E). \*\*\**\*P* < 0.0001 between Q175DN and WT; ^####^*P* < 0.0001 between Q175DN-SUMO1KO and Q175DN by Fisher’s exact test (for F), and \*\*\**\*P* < 0.0001 between Q175DN and WT; ^####^*P* < 0.001 between Q175DN-SUMO1KO and Q175DN by one-way ANOVA followed by Tukey’s multiple comparison test (for G).

### SUMO1 deletion abolishes striatal atrophy in HD mice

We also analyzed whether the behavioral protection induced by SUMO1 deletion is accompanied by changes in anatomical deficits. We sacrificed the mice after the behavioral analysis at approximately 13 months of age and examined striatal anatomical alterations by measuring the lateral ventricular area (an indirect measurement of striatal atrophy). We observed lateral ventricle enlargement in the Q175DN mice in the frontal, central, and caudal regions of the striatum, consistent with previous reports [97, 99] (Fig. 2A and B). This enlargement was markedly diminished in the Q175DN-SUMO1KO mice in all three striatal regions, and the lateral ventricle sizes were comparable to those of the WT and SUMO1KO control groups (Fig. 2A and B). Thus, SUMO1 deletion almost completely prevented the striatal atrophy observed in the knock-in HD mice.

**Figure 2.**
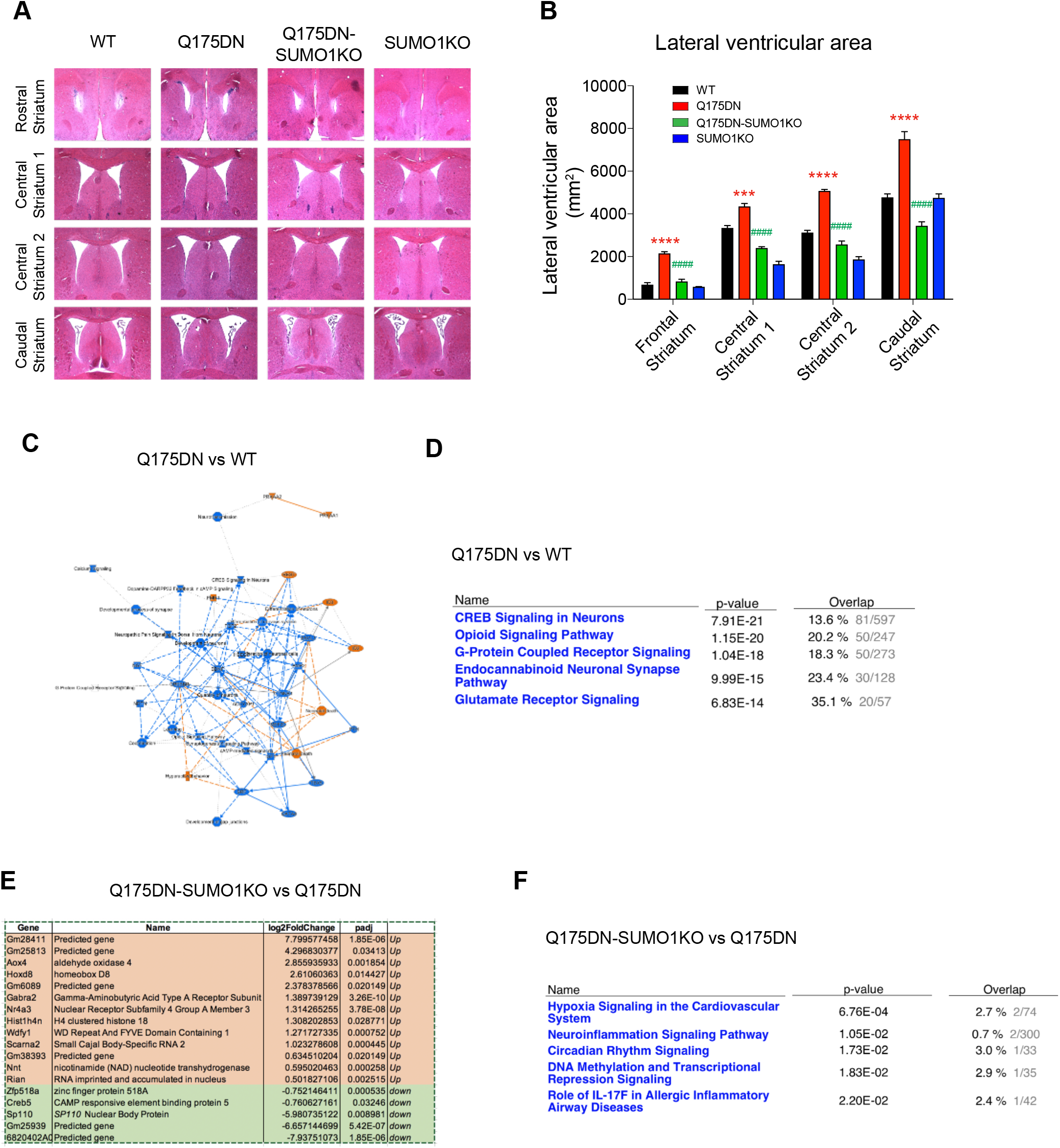
Effect of SUMO1 deletion on striatal anatomical changes and the striatal gene transcription in Q175DN mice. (A) Representative hematoxylin/eosin-stained sections for rostral (+1.45 from bregma), central striatum 1 (+1.0 from bregma), central striatum 2 (+0.60 from bregma), and caudal (+0.1 from bregma) lateral ventricles at the striatum level from the WT, Q175DN, Q175DN-SUMO1KO and SUMO1KO mice. (B) Quantification of the lateral ventricular area from A. Data are mean ± SEM (n = 5 mice per group, 3 – 4 sections were analyzed and averaged for each striatal region per mouse). \*\*\**P* < 0.001, \*\*\*\**P* < 0.0001 between Q175DN and WT; ^####^*P* < 0.0001 between Q175DN-SUMO1KO and Q175DN using two-way ANOVA followed by Tukey’s multiple comparison test (C, D) IPA network analysis between WT vs Q175DN groups. (E) List of differentially regulated genes in striatum from Q175DN-SUMO1KO mice compared to Q175DN mice. (F) IPA analysis between Q175DN and Q175DN-SUMO1KO.

### Effect of SUMO1 deletion on the striatal gene expression landscape in HD mice

Transcriptional dysregulation is a major event in HD pathogenesis [6, 7, 13, 101-103], and SUMO is involved in the transcriptional repression as well as activation functions [104-106]. We therefore examined whether SUMO1 impacts striatal atrophy in HD (Fig. 2A, B) by a dysregulation of gene expression. We performed a genome wide striatal tissue–specific gene expression pattern analysis by comparing RNA seq in the striatum of 13-month-old WT, Q175DN, Q175DN-SUMO1KO, and SUMO1KO mice (n = 3/group). Gene counts revealed 1042 genes that were significantly upregulated and downregulated (padj < 0.05) in the Q175DN compared to the WT striatum (Data file 1). Ingenuity pathway network analysis (IPA) revealed significant alterations in canonical pathways, such as cAMP-response element binding protein (CREB), opioid signaling, endocannabinoid signaling, G-protein-coupled receptors (GPCR), and glutamate receptor signaling, as well as fragile X mental retardation 1 (FMR1) signaling, between the WT and Q175DN striatum (Fig. 2C, D, Data file 1). Some of these altered pathways in Q175HD have been previously reported[107]. We found significant alterations of 255 genes between WT and SUMO1KO mice (padj < 0.05) and of 2308 genes between SUMO1KO and Q175DN mice (padj < 0.05); most of the altered genes involved opioid signaling, CREB signaling, and GPCR signaling, among others (Data file 1, Supp Fig. 1). However, we found very few differentially expressed genes between Q175DN-SUMO1KO and Q175DN mice striatum (padj < 0.05). Of the 18 genes discovered, 5 were downregulated and 13 were upregulated (Fig. 2E). Due to the small number of gene alterations, IPA analysis did not identify a comprehensive network of biological processes or pathways, as the number of connectible entities with sufficiently high z-scores was insufficient. However, IPA did hint that the modulated genes belonged to hypoxia, circadian rhythm, DNA methylation, and inflammatory processes (Fig. 2F) in Q175DN-SUMO1KO compared to Q175DN mice striatum. These results indicate that SUMO1 has only a nominal impact on the overall gene expression changes occurring in the HD striatum, suggesting that SUMO1 may promote striatal atrophy via post-transcriptional and/or post-translational mechanisms.

### SUMO1 deletion prevents the inflammatory responses of HD

Since the IPA analysis suggested that the linked altered genes were functionally enriched in inflammatory responses in Q175DN-SUMO1KO compared to Q175DN mice striatum (Fig. 2F), we carried out glial fibrillary acidic protein (GFAP) immunoreactivity assays to assess the inflammatory changes within the frontal, central, and caudal regions of striatal sections from the WT, Q175DN, Q175DN-SUMO1KO, and SUMO1KO mice (Fig. 3A). GFAP expression was strikingly enhanced in all regions, with a slightly increased signal in the central striatum, in Q175DN mice compared to WT mice (Fig. 3A, B), consistent with previous reports [108, 109]. The Q175DN-SUMO1KO mice showed a marked decrease in GFAP immunoreactivity in all three regions compared to the Q175DN mice (inset c and e), whereas GFAP immunoreactivity was unaltered between the WT and SUMO1KO striatum (Fig. 3A, B). Higher GFAP expression is also seen in the cortical region of Q175DN mice (inset d) which is also diminished in the Q175DN-SUMO1KO (inset f).

**Figure 3.**
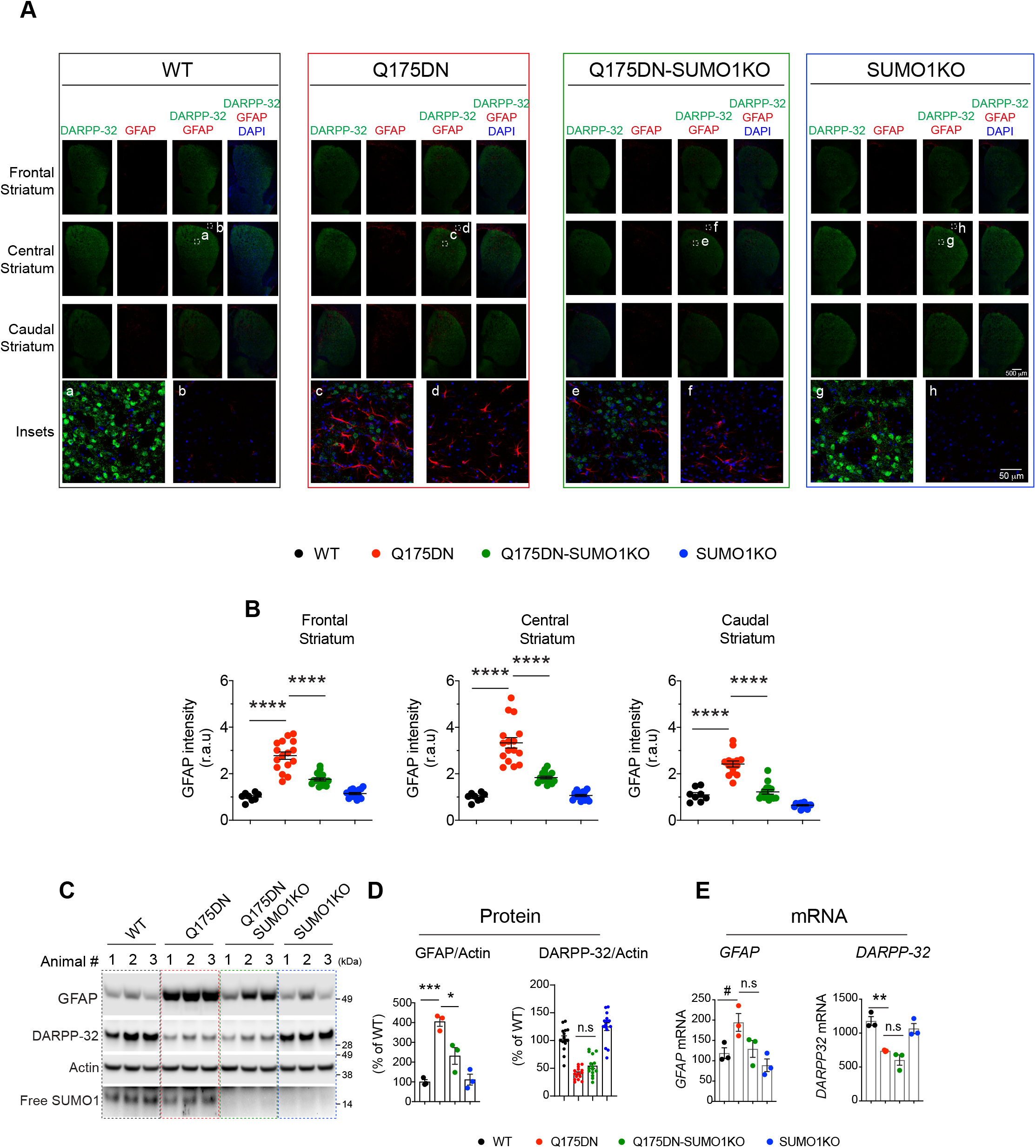
SUMO1 deletion diminishes inflammatory astroglial response in Q175DN mice. (A) Representative confocal images for the frontal, central and caudal striatum showing DARPP-32 (green), and GFAP (red) IHC, and nuclear stain DAPI (blue) with the indicated magnified image in the insets from the WT, Q175DN, Q175DN-SUMO1KO and SUMO1KO mice. Insets, striatum (a, c, e, g) and cortex (b, d, f, h). (B) Quantification of the GFAP intensity from the frontal, central and caudal striatum. Data are mean ± SEM, n = 5 mice per group, between 1 to 3 sections for each striatal region per mouse were analyzed (total 8 sections for WT, and 16 sections each for Q175DN, Q175DN-SUMO1KO and SUMO1KO groups). \*\*\**\*P* < 0.0001 by one-way ANOVA followed by Tukey’s multiple comparison test. (C) Western blots for the GFAP, DARPP-32 and Free SUMO1 expression for all the genotypes. (D) Quantification of the blots from C. Data are mean ± SEM, n = 3 mice per group for GFAP, n = 3 mice per group for DARPP-32. *\*P* < 0.05, and \*\*\**P* < 0.001 by one-way ANOVA followed by Tukey’s multiple comparison test. (E) TPM (Transcripts Per Kilobase Million) counts of *GFAP* and *DARPP-32* mRNA from the RNA-seq from striatum of WT, Q175DN, Q175DN-SUMO1KO and SUMO1KO mice. Data are mean ± SEM, n = 3 mice per group. *\*\*P* < 0.01 by one-way ANOVA followed by Tukey’s multiple comparison test. ^*#*^*P* < 0.05 by unpaired two-tailed Student’s *t* test.

Western blot analysis of the striatal tissue further confirmed that GFAP protein expression, but not mRNA expression, was significantly reduced in the Q175DN-SUMO1KO mice compared to the Q175DN mice (Fig. 3C, D, E). The protein levels of DARPP-32, a commonly used biochemical MSN marker for striatal damage in HD [110, 111], was diminished in Q175DN mice compared to WT mice. This reduction is likely due to diminished DARPP-32 mRNA in Q175DN mice (Fig. 3E). The Q175DN-SUMO1KO mice showed a similar extent of loss of DARRP-32 protein and mRNA, indicating SUMO1 may not interfere with the transcriptional downregulation of DARPP-32 in the HD striatum (Fig. 3E). These results indicate that SUMO1 deletion prevents the striatal atrophy that is accompanied by a strong reduction in the inflammatory glial response in the striatum.

### SUMO1 deletion enhances EM48+ aggregates and decreases soluble forms of HTT

As SUMO plays a major role in regulating protein aggregation, including aggregation of mHTT [60, 84, 86, 112-115], we also analyzed the aggregated and soluble forms of mHTT by immunohistochemistry (IHC) and western blotting. We used EM48 antibody, the most sensitive antibody for detecting mHTT inclusions [116, 117], as well as MAB2166 (clone 1HU-4C8) [117, 118] and polyQ (clone 3B5H10) [119, 120] antibodies to detect FL-HTT. Immunofluorescent staining of striatal brain sections of WT or SUMO1KO mice with EM48 antibody did not reveal any inclusions (data not shown), so these mice were not included in further analysis. However, EM48 antibody readily detected HTT aggregates in Q175DN and Q175DN-SUMO1KO striatal brain sections in both the cortex and striatum (Fig. 4A), consistent with previous report [97]. The numbers of DARPP-32+ neurons between Q175DN and Q175DN-SUMO1KO mice did not differ (Fig. 4A, B); however, a stronger colocalization of EM48+ punctae was observed in DARPP-32 MSNs (Fig. 4C, D) in the striatum of Q175DN-SUMO1KO mice than in Q175DN mice. These data indicated that SUMO1 deletion altered the mHTT distribution in MSNs. Therefore, we further analyzed the distribution of EM48+ aggregates in the cell soma, nucleus, and extracellular space.

Confocal analysis revealed no differences in the numbers of cells (DAPI+) or total EM48+ mHTT aggregates between Q175DN and Q175DN-SUMO1KO mice (Fig. 4E, F, G). However, the number of nuclear EM48+ aggregates (solid arrowhead, Fig. 4E, H) in the DARPP-32+ neurons and extracellular EM48+ aggregates (asterisk) was significantly diminished (Fig. 4E, I), whereas the cytoplasmic EM48+ aggregates (open arrowhead) in DARPP-32+ neurons were increased in the Q175DN-SUMO1KO mice compared to the Q175DN mice (Fig. 4E, J).

**Figure 4.**
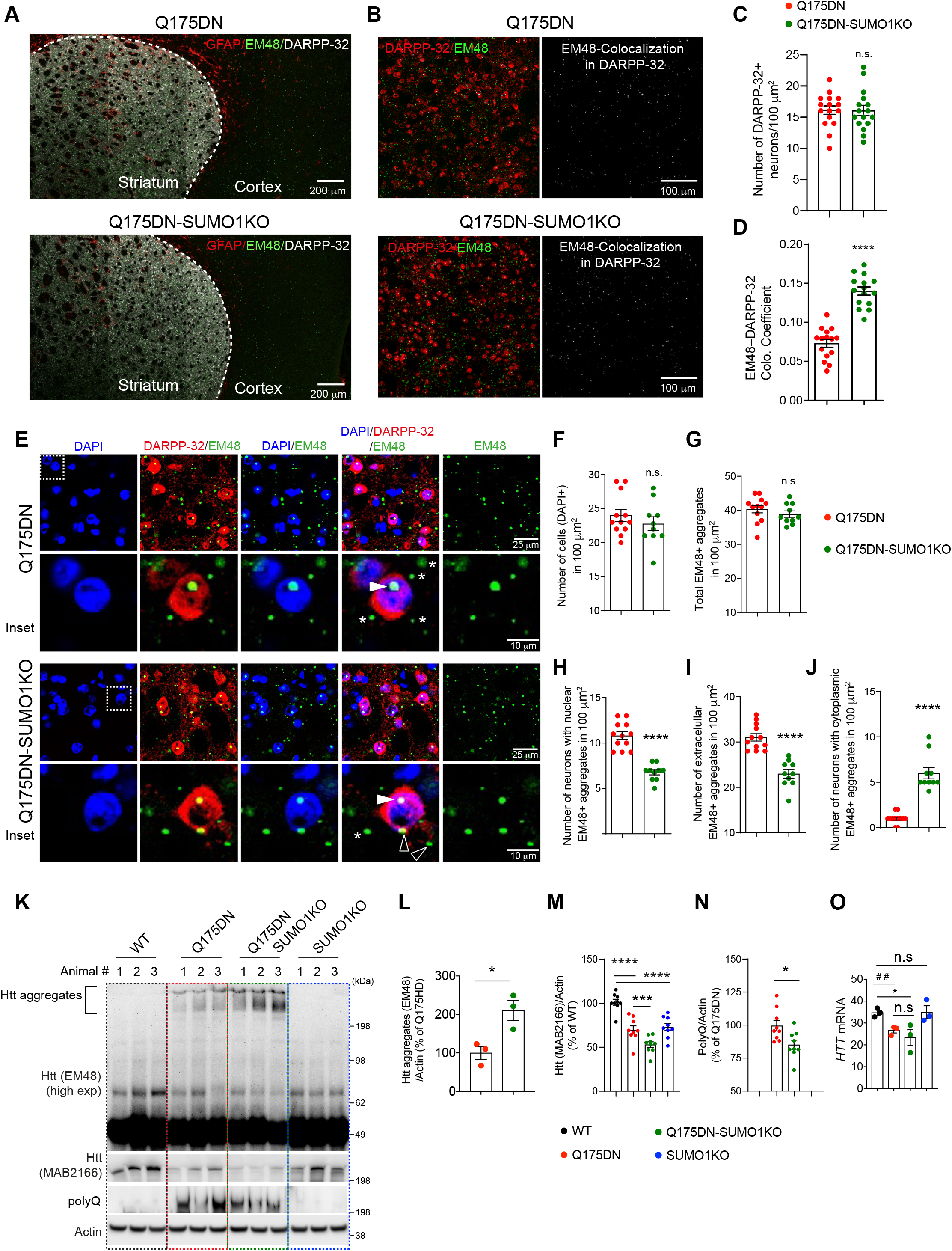
SUMO1 deletion alters EM48 aggregate distribution in the Q175DN mice striatum. (A) Low magnification confocal immunofluorescence images showing GFAP (red), Htt aggregates (EM48+, green), and DARPP-32 (white) expression from Q175DN and Q175-SUMO1KO brain sections. (B) Representative confocal images showing EM48 aggregates (green) and DARPP-32 (red) IHC and colocalization of EM48 aggregates and DARPP-32 in the striatum from Q175DN and Q175-SUMO1KO mice. (C) Bar graphs show the number of DARPP-32+ neurons in 100µm^2^ area. Data are mean ± SEM (n = 5 mice per group, from each mouse 4 areas were averaged from 2-4 sections and section averages were used for analysis). n.s. not significant by two-tailed unpaired Student’s t-test. (D) Mander’s overlay coefficient of EM48 and DARPP-32 from B. Data are mean ± SEM (n = 5 mice per group, from each mouse 4 areas were averaged from 2-4 sections and section averages were used for analysis). \*\*\*\**P <* 0.0001 by two-tailed unpaired Student’s t-test. (E) Confocal images showing the EM48 aggregates (green), and DARPP-32 (red) IHC, and DAPI. Lower panels show the magnified insets depicting nuclear EM48+ aggregates (solid arrowhead), cytoplasmic EM48+ aggregates (open arrowhead) in the DARPP-32+ neuron, and extracellular EM48+ aggregates (asterisk). (F-J) Quantifications for the total number of cells (DAPI+) (F), total EM48+ aggregates (G), number of neurons with nuclear EM48+ aggregates (H), number of extracellular EM48+ aggregates (I) number of neurons with cytoplasmic aggregates (J) in 100µm^2^ area. Data are mean ± SEM (n = 5 mice per group, from each mouse 4 areas were averaged from 2-4 sections and section averages were used for analysis). \*\*\*\**P <* 0.001, n.s. not significant by two-tailed unpaired Student’s t-test. (K) Represntative western blot for indicated proteins from the striatum of WT, Q175DN, Q175DN-SUMO1KO and SUMO1KO mice. (L-N) Quantification of the EM48 aggregates (L), Htt levels (MAB2166) (M) and PolyQ (N) from K. Data are mean ± SEM (n = 3-9 mice per genotype). \**P <* 0.05 by two-tailed unpaired Student’s t-test; \*\*\**P <* 0.001, \*\*\*\**P <* 0.0001, by one-way ANOVA followed by Tukey’s multiple comparison test. (O) TPM (Transcripts Per Kilobase Million) counts of *Htt* RNA from the RNA-seq from striatum of WT, Q175DN, Q175DN-SUMO1KO and SUMO1KO mice. Data are mean ± SEM (n = 3 independent experiments). \**P <* 0.05, by one-way ANOVA followed by Tukey’s multiple comparison test. ^##^*P* < 0.01 by two-tailed unpaired Student’s t-test.

Western blotting analysis of striatal brain lysates prepared in non-denaturing Tris buffer (which enriches cytoplasmic proteins [121, 122]) showed a significant enhancement of EM48+ aggregates in the Q175DN-SUMO1KO striatum, compared to the Q175DN striatum (Fig. 4K, L), supporting confocal findings that SUMO1 deletion increased the cytoplasmic EM48+ mHTT levels. As expected no EM48+ aggregates were seen in WT and SUMO1KO control groups (Fig. 4K). We then probed the same lysates for MAB2166 antibody (which detects both HTT and mHTT) or polyQ antibody to detect full length (FL)-mHTT and mHTT. We found diminished levels of HTT levels as well as poly Q mHTT in the Q175DN mice striatum, consistent with earlier reports [123, 124], but a further diminishment in the Q175DN-SUMO1KO striatum (Fig. 4K, M). Note, normal HTT protein levels are also diminished in SUMO1KO striatum (Fig. 4K, M). Next we determined if SUMO1 regulates the *HTT* mRNA. We found that while *HTT* mRNA is significantly diminished in the Q175DN mice compared to WT, it remained diminished to a similar extent in the Q175DN-SUMO1KO mice. The mRNA of normal *HTT* is similar between WT and SUMO1KO striatum (Fig. 4K, O). Thus, SUMO1 deletion reduces the protein levels of mHTT as well as normal HTT, but not their mRNA levels, in the striatum, indicating post-transcriptional and/or post-translational mechanisms.

### SUMO1 deletion enhances the p62 and EM48+ interaction and autophagic activities in the striatum

Next we explored possible mechanisms for the regulation of HTT/mHTT levels by SUMO1 in the striatum. We focused on p62, a selective macro-autophagy receptor that sequesters ubiquitinylated proteins and regulates protein aggregation [125]. Moreover, previous studies have identified p62 as a critical mediator of HTT-induced selective autophagy and disease phenotype [30, 31, 120, 126-130]. We carried out p62 immunostaining in brain sections. By applying fluorescence colocalization analysis using Manders overlap coefficient (MOC) [131, 132] we found that p62 strongly associated with EM48+ aggregates in Q175DN striatum and the association was significantly diminished in the Q175DN-SUMO1KO striatum (arrowhead, Fig. 5A, B).

**Figure 5.**
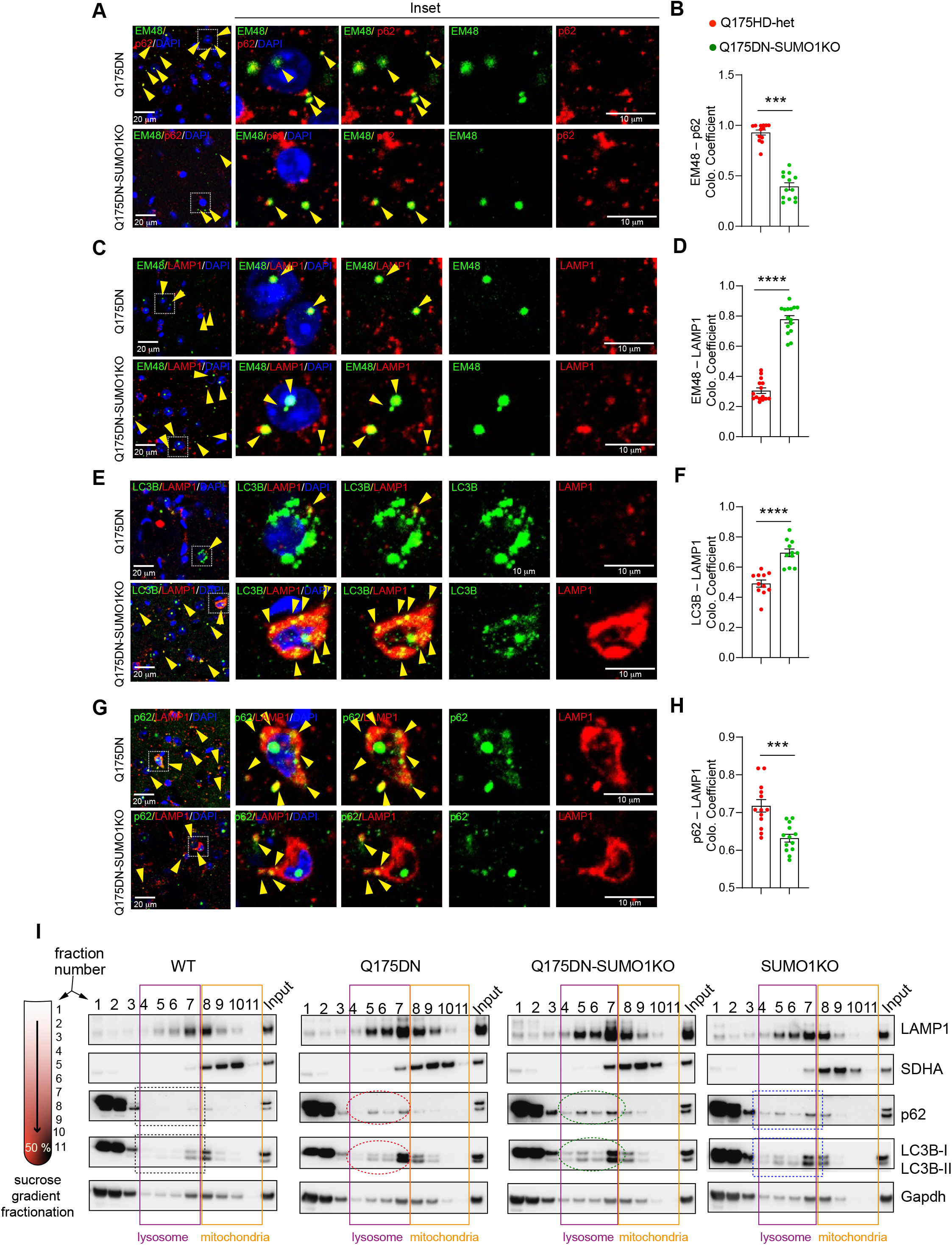
SUMO1 deletion enhances autophagic activities in the striatum. (A-H) IHC and quantification for colocalization coefficients for EM48 – p62 (A, B), EM48 – LAMP1 (C, D), LC3B – LAMP1 (E, F) and p62 – LAMP 1 (G, H) in the striatum from Q175DN and Q175-SUMO1KO mice. DAPI (blue) and color channels (green or red) according to indicated antibody combination. Insets with magnified images is shown on the right. Yellow arrowheads indicate the colocalization for green and red channels. Data are mean ± SEM (n = 5 mice per group, from each mouse 4 areas were averaged from 2-4 sections and section averages were used for analysis). \*\*\**P <* 0.001, \*\*\*\**P <* 0.0001 by two-tailed unpaired Student’s t-test. (I) Western blot of the biochemical fractionation from striatal tissues of indicated genotypes. Purple and orange rectangles show lysosomal and mitochondrial fractions, respectively. Dotted rectangle and oval show enrichment of indicated proteins.

Simultaneous binding occurs between p62 and autophagosomes via the interaction with LC3B-II and between p62 and mHTT/HTT [133, 134]. We used the MOC to examine the association of EM48+ mHTT and LAMP1 (a lysosomal marker), the association between LC3B-II (an autophagosome marker) and LAMP1, and the p62 and LAMP1 co-localizations. We found strong increases in the EM48+ association with LAMP1 (arrowhead, Fig. 5C, D), as well as enhanced interaction between LC3B-II and LAMP1 (arrowhead, Fig. 5E, F). By contrast, the p62 and LAMP1 association was diminished in the Q175DN-SUMO1KO striatum compared to the Q175DN striatum (arrowhead, Fig. 5G, H).

These confocal microscopy results suggested that SUMO1 deletion may enhance autophagic activities in the HD striatum. We then separated lysosomes using sucrose density gradient centrifugation and found enhanced p62 and LC3B-II accumulation in the lysosomal fractions in the Q175DN-SUMO1KO (green oval) compared to the Q175DN striatum (red oval) (Fig. 5I). The SUMO1KO mice also displayed enhanced p62 and LC3-II accumulation (blue rectangle) in the lysosomal fraction compared to the WT (black rectangle).

Collectively, these results indicate that SUMO1 deletion enhances autophagy activities in the HD striatum.

### SUMO1 deletion increases autophagy flux in a cellular model of HD

We next used cultured HD striatal cells to examine the cause-and-effect relationship between SUMO1 and autophagy regulation. We depleted SUMO1 by CRISPR/Cas-9 in well-established *STHdh*^Q7/Q7^ (WT control) and *STHdh*^Q7/Q111^ (HD-het) cell lines [135] and successfully depleted ∼50–60% of the SUMO1 modifications (Fig. 6A, B). The steady state levels of HTT were not significantly affected by diminished SUMOylation (Fig. 6A, B). The HD-het cells showed enhanced basal LC3B-II compared to WT control and the levels were not affected by SUMO1 depletion (Fig. 6A, B). By contrast, enhancement of p62 in HD-het cells was diminished by SUMO1 depletion (Fig. 6A, B). Confocal analysis of SUMO1-depleted (SUMO1Δ) control and HD-het cells showed that colocalization (MOC) of EM48 – p62 (Fig. 6 C, D), LAMP1 – p62 (Fig. 6 E, F), and EM48 – LAMP – p62 (Fig. 6 G, H) were decreased in SUMO1Δ HD-het cells compared to SUMO1 intact HD-het (control) cells. These results suggest that autophagy activities are increased upon SUMO1 deletion in HD cells.

**Figure 6.**
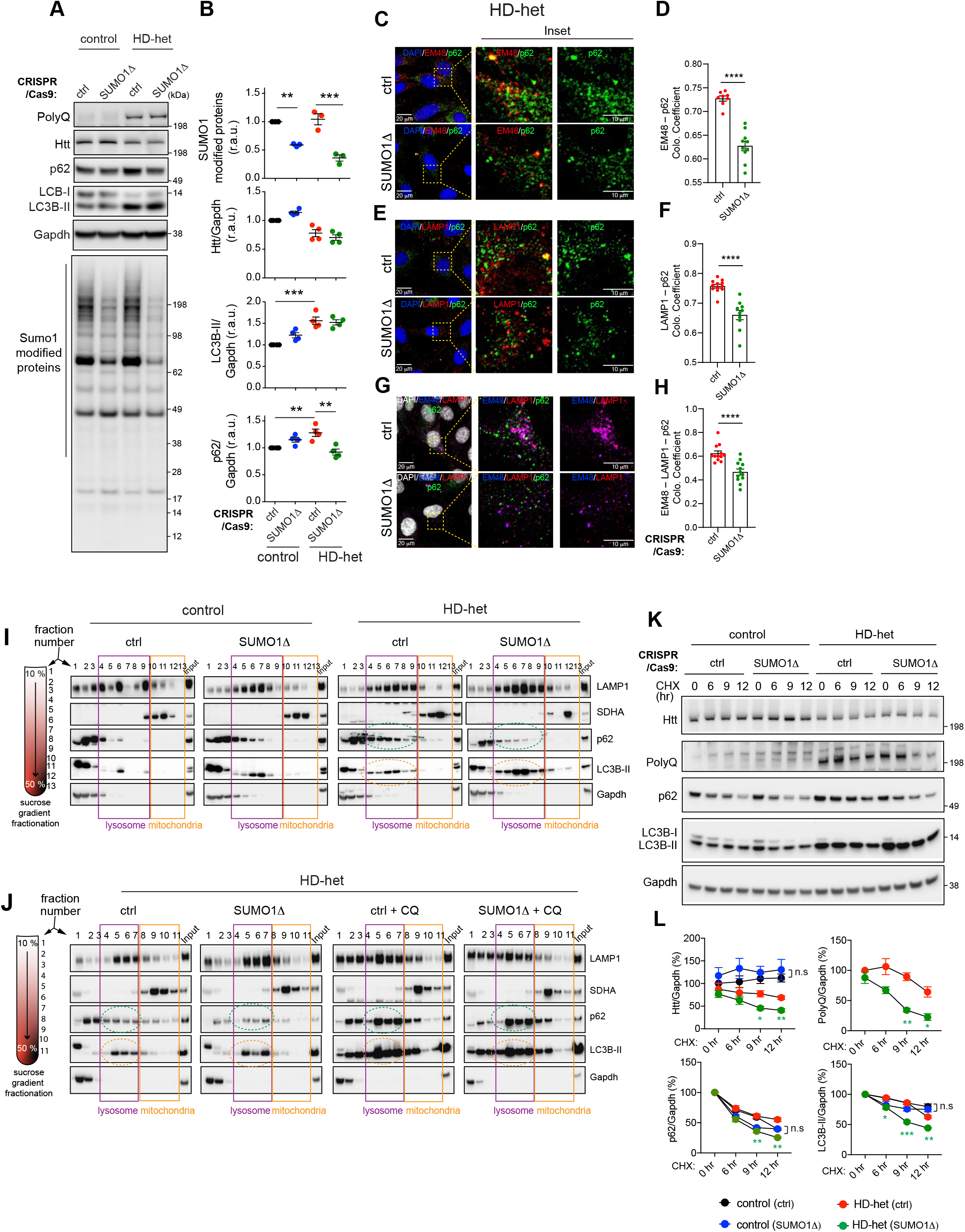
SUMO1 deletion increases the autophagy in a cellular model of HD. (A). Western blotting of the indicated proteins in control (ctrl) and SUMO1 deleted (SUMOΔ) control and HD-het striatal cells. (B) Quantification of the indicated proteins from A. Data are mean ± SEM (n = 3 – 4 independent experiments). \*\**P <* 0.01, \*\*\**P <* 0.001, by one-way ANOVA followed by Tukey’s multiple comparison test. (C-H) Confocal images and corresponding magnified insets of control (ctrl) and SUMO1 deleted (SUMOΔ)-HD-het cells, immunostained and showing quantification of Mander’s colocalization coefficient for EM 48 – p62 (C, D), LAMP1 – p62 (E, F) and EM48 – LAMP1 – p62 (G, H). Data are mean ± SEM (n = 4 independent experiments, from each experiment two–three100 µm^2^ areas were quantified and used for group analysis). \*\*\*\**P <* 0.0001 by two-tailed unpaired Student’s t-test. (I, J) Western blot for the biochemical fractionation assay from control (ctrl) and SUMO1 deleted (SUMOΔ) cells and without (I) or with chloroquine, CQ 50 µM treatment in HD-het cells. Purple and orange rectangles show lysosomal and mitochondrial fractions, respectively. Dotted oval show enrichment of indicated proteins. Representative western blot of indicated proteins from ctrl and SUMOΔ cells with cycloheximide (CHX, 75 µM) treatment for the indicated time points. (L) Quantification of indicated proteins from K, normalized with GAPDH. Data presented as mean ± SEM (n = 4 independent experiments), \**P* < 0.05, \*\**P* < 0.01, and \*\*\**P* < 0.001 HD-het (SUMOΔ) vs HD-het (ctrl) cells, by repeated measure two-way ANOVA followed by Tukey’s multiple comparison test or Bonferroni’s multiple comparisons test (for PolyQ). n.s: not significant.

Biochemical fractionation experiments further confirmed an increased autophagic activity, namely the high LC3B-II association (orange circle) and the diminished p62 levels (green circle) in the lysosomal compartment, in SUMO1Δ HD-het cells compared to ctrl HD-het cells (Fig. 6I). Assessment of autophagy flux with chloroquine showed further accumulation of LC3B-II and p62 in the SUMO1Δ HD-het cells (Fig. 6J), suggesting that SUMO1 deletion most likely increases autophagy flux in HD.

Unlike the in vivo paradigm, we did not observe any drastic decline in basal HTT levels in SUMO1Δ HD-het cultured cells (Fig. 6A, B). One reason could be that steady-state effect of SUMO1 on mHTT degradation by autophagy cannot be distinguishable from its synthesis. So, we carried out kinetics experiments with protein synthesis inhibitor cycloheximide (CHX) and found that both mHTT and poly-Q mHTT levels were strongly diminished in the presence of CHX in the SUMO1Δ HD-het cells compared to the WT control (Fig. 6K, L). We also observed more autophagic activity evident by increasesd LC3B-II expression and decreased p62 in the SUMO1Δ HD-het cells (Fig. 6K, L). We did not detect any change in the normal HTT levels, further indicating that in vivo regulation of HTT by SUMO1 is perhaps regulated in a tissue- and age-dependent manner (Fig. 6K, L). These results indicate SUMO1 deletion diminishes mHTT levels by enhancing autophagic activities.

### The SUMOylation inhibitor ginkgolic acid (GA) enhances autophagy in neuronal HD cells and human HD fibroblasts

We then examined whether the pharmacological modulation of SUMOylation affected autophagy activities in HD cells. To determine this, we investigated the effect of ginkgolic acid (GA). GA, an extract from *Ginkgo biloba* leaves, is a type of alkyl phenol that can directly bind SUMO-activating enzyme, E1, and inhibit the formation of the E1-SUMO intermediate, thereby diminishing SUMOylation [136-140]. Because SUMO1 depletion enhanced autophagic activities in HD, we hypothesized that GA might enhance the autophagy activities and alter HTT levels in HD cells. We tested the GA effect in WT control, HD-het striatal neuronal cells and human unaffected with 17CAG repeats and HD-het fibroblasts that contain mHTT with 69 CAG repeats (this is closer to the form of mHTT in juvenile HD).

Treatment with GA (100 μM) did not cause observable toxicity but resulted two major effects. First, at 0 hr, GA induced a remarkable upregulation of LC3B-II and p62 expression in striatal cells (Fig. 7A, B) as well as fibroblasts (Fig. 7C, D). Second, upon time course experiments with CHX, the GA treatment did not affect normal HTT or mTOR levels both in WT striatal neuronal cells (Fig. 7A, B) and unaffected fibroblasts (Fig. 7C, D). However, GA showed a strong trend of diminished mHTT levels, particularly evident with poly-Q antibodies in both cell types, and the diminishing effect of GA appeared more pronounced in human HD fibroblast (Fig. 7B, D). Moreover, in CHX treatment, we observed time-dependent rapid degradation of autophagic cargo, p62, in HD striatal cells and human fibroblast cells in presence of GA compared to vehicle treated controls. This data clearly indicate SUMOylation inhibitor GA promotes robust autophagy and rapid clearance of autophagy cargo. The effect of GA is specific to mHTT because GA did not influence normal HTT or mTOR levels compared to vehicle controls (Fig. 7A-D). These results indicate that GA mimics SUMO1 depletion and potentiates autophagy activities and diminishes mHTT protein levels in both mouse and human HD cells.

**Figure 7.**
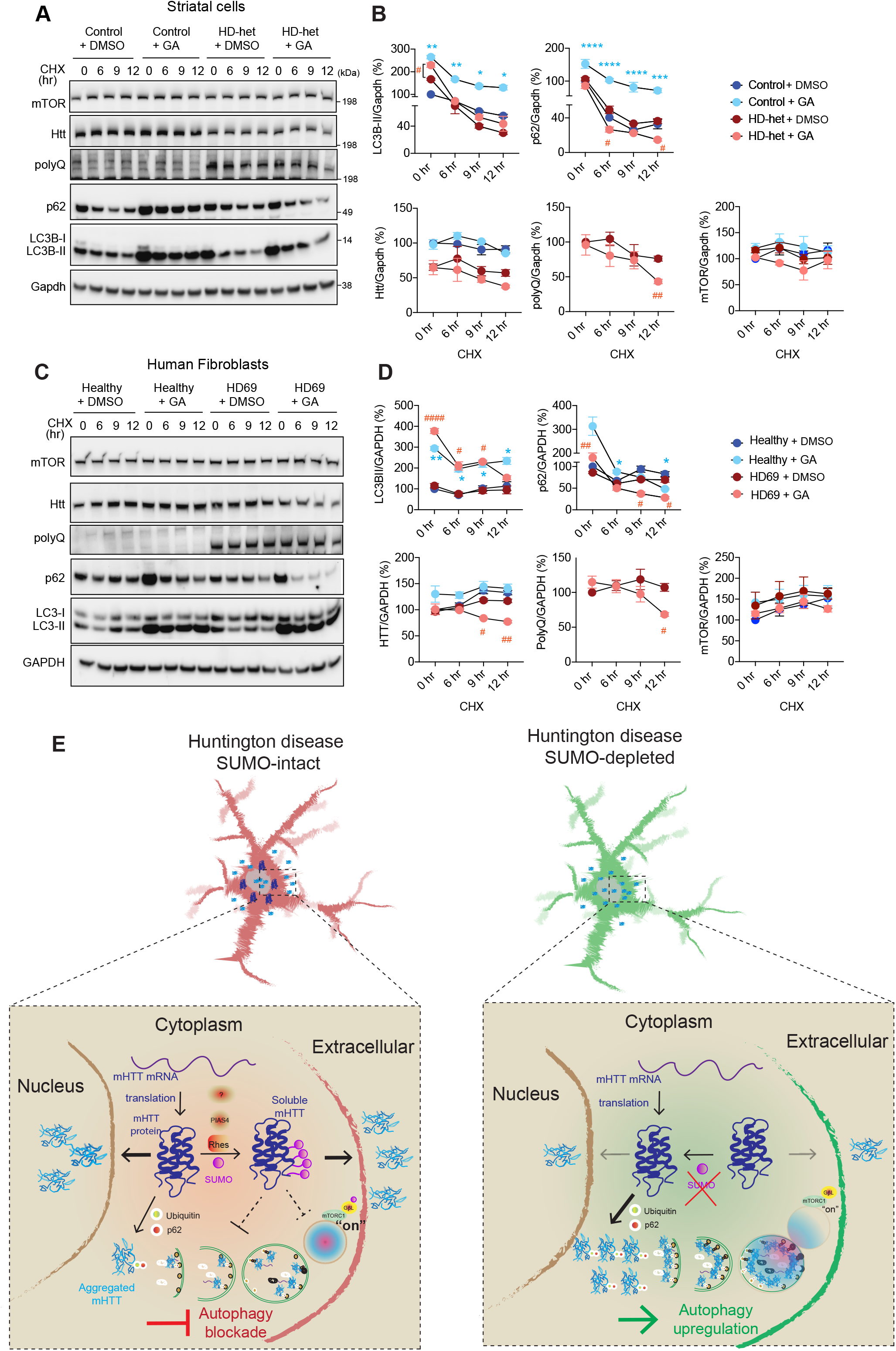
SUMO inhibition by Ginkgolic acid (GA) decreases stead-state-levels of mHTT in HD striatal and human HD fibroblasts. (A) Representative western blot for the indicated proteins from control and HD-het cells treated with vehicle (DMSO) or Ginkgolic acid (GA, 100 µM) for 24 hr and CHX (75 µM) for indicated time points. (B) Quantification of indicated proteins from A, normalized with GAPDH (vehicle treated control cells as 100%). Data presented as mean ± SEM (n = 3 independent experiments), \**P* < 0.05, \*\**P* < 0.01, \*\*\**P* < 0.001, and \*\*\*\**P* < 0.0001 between control and control + GA; ^#^*P* < 0.05, and ^##^*P* < 0.01 between HD-het and HD-het + GA by repeated measure two-way ANOVA followed by Tukey’s multiple comparison test or Bonferroni’s multiple comparisons test (for PolyQ). (C) Human fibroblasts healthy or patient-derived HD cells (HD 69) treated with GA (100 µM) for 24 hr and CHX (50 µM) for indicated time points. Data presented as mean ± SEM (n = 3 independent experiments), \**P* < 0.05, and \*\**P* < 0.01 between healthy and healthy + GA; ^#^*P* < 0.05, ^##^*P* < 0.01, and ^####^*P* < 0.0001 between HD69 and HD69 + GA by repeated measure two-way ANOVA followed by Tukey’s multiple comparison test or Bonferroni’s multiple comparisons test (for PolyQ). **(E) A mechanistic model depicting SUMO1-mediated striatal abnormalities in HD**. *Left panel*. A properly folded mHTT protein is modified by SUMO1 keeping it as an enhanced state of solubility. Non SUMOylated, improperly folded mHTT can aggregate, which are capable of being degraded by autophagy or another degradation system. But SUMOylated mHTT may block autophagy by various means either by upregulating mTORC1 or by blocking one or more other steps in the autophagy assembly. The presence of SUMO1 may also enhance the export of mHTT to the nucleus and to the extracellular space, which may eventually get aggregated but cannot be degraded due to lack of degradation machinery. *Right panel*. Lack of SUMO1 relieves inhibitory effect on autophagy. More mHTT gets aggregated in the cytoplasm and are degraded via autophagy and other degradation pathway. This process diminishes the transport of mHTT to the nucleus as well as extracellular space. Collectively, mHTT toxic effects are eliminated upon deletion of SUMO1.

## DISCUSSION

Although polyglutamine expansion is sufficient for pathogenesis, the toxicity of mHTT is altered by additional cis/trans regulators. Previous studies have revealed that SUMO is a trans-acting regulator of mHTT toxicity in HD cells and in HD animal models [83, 84, 86, 141]. SUMO modification of mHTT has been shown previously to increase the soluble forms and to promote cellular toxicity [142-144], but no in vivo role or underlying mechanism for SUMOylation has been proposed for HD pathogenesis in knock-in model.

The present data reveal that SUMO1 deletion in the Q175DN HD-het knock-in mice diminished HD-related behavioral and anatomical deficits and inflammatory responses and was accompanied by the enhancement of EM48+/p62 localization, autophagy, and lysosomal activities in the striatum. Interestingly, the SUMO1-deleted MSNs showed alteration of mHTT from a soluble to an aggregated state, and the aggregates were then further reallocated from the nucleus and extracellular space to the cytoplasm. Thus, SUMO1 regulation of HD pathogenesis is accompanied by altered autophagy signaling and a potentially neurotoxic localization of soluble forms of mHTT in the cytoplasm. Notably, only a limited number of gene transcripts and the associated inflammatory biomarkers were altered in the striatum of SUMO1-deleted HD mice. These results reveal that the possible mechanism(s) may involve post-transcriptional regulation of SUMO signaling and an important autophagy regulatory mechanism in the mediation of mHTT-induced striatal atrophy.

Previous studies have linked SUMOylation to autophagy. For example, SUMOylation of Vps34 and its interaction with Beclin-1 is implicated in autophagosome formation in cancer and smooth muscle cells [76, 145]. Similarly, the SUMO E2 ligase Ubc9 can regulate autophagy flux in cardiomyocytes[146]. Upon DNA damage, SUMOylation of RhoB promotes its interaction with TSC2 and translocation to the lysosomes, where it inhibits mTORC1 to result in activation of autophagy [147]. Furthermore, deletion of the *Senp3* gene that codes for a de-sumoylating enzyme increased autophagy flux in the liver [148]. These results suggest that SUMOylation promotes autophagy flux. By contrast, blocking the SUMO pathway by depleting SUMO1 and UBC9 or by exposure to ginkgolic acid (a SUMOylation inhibitor) enhances the autophagy flux in breast cancer cells [149]. Thus, SUMOylation can modulate autophagy in two opposite directions, depending on the cell/tissue type, regulator, and targets of the SUMO pathway being altered. Therefore, a highly complex and intricate regulation of SUMOylation is associated with autophagy signaling.

Our results indicate SUMO1 in HD inhibits autophagy and that the mechanisms may involve a combination of multiple molecular processes. For example, SUMO1 modification of mHTT may directly interfere with the autophagy components; consequently, a loss of SUMO1 increases the levels of the ubiquitinated version of mHTT that, in turn, can interact strongly with p62 to enhance macroautophagy as well as mHTT clearance (Fig. 7E). This possibility is supported by our results showing enhanced autophagy activities and degradation of soluble mHTT in SUMO1-deleted HD striatum and cell models. SUMO may also inhibit autophagy via an indirect and integrated inhibitor feedback loop. For instance, 1) SUMO modification of GβL, which is a major component of mTOR, regulates nutrient-induced mTORC1 signaling, which inhibits autophagy [18, 150]. 2) SUMO modification of mHTT, which also promotes nutrient-induced mTORC1 signaling[100], may further inhibit autophagy [150]. And 3) SUMOylation of Rhes on multiple sites may regulate Rhes-mediated Beclin-1-dependent and mTORC1-independent autophagy [82, 151]. Thus, SUMO deletion may regulate autophagy in the striatum via both mTORC1-dependent and mTORC1-independent signaling pathways.

We propose a model (Fig. 7E) whereby mHTT is synthesized and then undergoes SUMO modification, mediated by one or more SUMO E3-like proteins, such as Rhes [81, 82, 86], PIAS [83, 141], or unknown E3s. Both the non-SUMOylated and SUMOylated versions of mHTT can move into the nucleus and out of the neurons (into the extracellular space) and eventually become aggregated. A portion of the non-SUMOylated HTT that aggregates in the cytoplasm is removed by the ubiquitin and p62 interaction through autophagy or other degradation pathways, such as proteasomes [152, 153]. By contrast, the SUMOylated soluble mHTT forms inhibit autophagy, either by activating mTORC1 or blocking autophagosome formation or lysosomal activities [134, 154, 155]. Thus, SUMO1 causes a slow and sustained blockade of autophagy, thereby affecting neuronal homeostasis and leading to the MSN dysfunction observed in HD (Fig. 7E, left panel). Lack of SUMO1 averts this autophagy inhibition and impedes the transport of mHTT to the nucleus and extracellular space. This enhances the formation of mHTT aggregates in the cytoplasm, and these can now be degraded through the active autophagy pathway (Fig. 7E, right panel).

Surprisingly, the SUMO E3-like protein Rhes levels are downregulated in the Q175DN HD and Hdh150Q knock-in mouse striatum (data not shown), indicating that compensatory mechanisms are at work. Perhaps clearance of Rhes will decrease the SUMOylated forms of mHTT and extend the onset and reduce the severity of the disease process. Consistent with this notion, forced expression of Rhes enhanced soluble forms of mHTT and worsened the motor phenotype in HD mouse models[92].

Overall, this report, based on data integrated from in vivo models, behavioral approaches, and biochemical tools, provides insights into a critical role of SUMO in autophagy regulation. SUMO1 deletion reduces the soluble mHTT levels, prevents the inflammatory response, and ameliorates HD pathogenesis. This research shows that interfering with SUMO signaling for example by using pharmacological inhibitor such as GA or its analogs can provide therapeutic opportunities to relieve HD symptoms and progression by upregulating autophagy and diminishing toxic mHTT levels.

## MATERIALS AND METHODS

### Reagents and antibodies

All reagents were purchased from Millipore-Sigma unless indicated otherwise. The following commercial antibodies were used: Huntingtin (WB-1:3000 clone 1HU-4C8, no. MAB2166), EM48 huntingtin (WB-1:10000, IHC-1:250, no. MAB5374), and Anti-polyglutamine (poly-Q) antibody (WB-1:5000, no. P1874) antibodies were obtained from Millipore-Sigma. Antibody against GFAP (IHC-1:500, 13-0300) was from ThermoFisher Scientific and LAMP1 (WB-1:1000, 1:200 (IHC), no. 1D4B) was from Developmental Studies Hybridoma Bank. Antibodies for DARPP-32 (WB-1:20000, IHC-1:500, no. 2306), SUMO1 (WB-1:1000, no. 4930), p62 (WB-1:1000, no. 39749), p62 (WB-1:1000, IHC-1:200, no. 23214), p62 (WB-1:1000, no. 7695), SDHA (WB-1:1000, no. 11998), LC3B (WB-1:2000, IHC-1:200, no. 3868), and mTOR (WB-1:3000, no. 2983) were from Cell Signaling Technology. Antibodies for β-actin (WB-1:20,000; no. sc-47778), glyceraldehyde-3-phosphate dehydrogenase (Gapdh) (WB-1:1000; no. sc-32233) were obtained from Santa Cruz Biotechnology. HRP-conjugated secondary antibodies: goat anti-mouse (1:10,000; no.115-035-146, goat anti-rabbit (1:10,000; no. 111-035-144) and goat anti-rat (1:10,000; no. 112-035-143) were from Jackson ImmunoResearch Inc. Donkey anti-Mouse Alexa Fluor 488 (1:500, no. A21202), Donkey anti-Mouse Alexa Fluor 568 (1:500, no. A10037), Donkey anti-Mouse Alexa Fluor 647 (1:500, no. A31571), Donkey anti-Rabbit Alexa Fluor 488 (1:500, no. A21206), Donkey anti-Rabbit Alexa Fluor 568 (1:500, no. A10042), Donkey anti-Mouse Alexa Fluor 647 (1:500, no. A31573), Donkey anti-Mouse Alexa Fluor 568 (1:500, no. A10037), and Donkey anti-Rat Alexa Fluor 594 (1:500, no. A21209)–conjugated secondary antibodies were from ThermoFisher Scientific.

### Animals

*Sumo1* knock out mice (SUMO1KO) were obtained from Jorma Palvimo [78]. zQ175 neo-deleted knock-in heterozygous (Q175DN HD-het, #029928) and C57BL/6J control mice were from Jackson laboratories. Q175DN (HD-het) and SUMO1KO mice were bred to produce Q175DN-SUMO1KO and littermate groups, Q175DN (Het), SUMO1KO and WT, and the genotypes were confirmed by an established PCR protocol. Behavior evaluation was performed as indicated in the timeline Figure 1A. Number of animals used are indicated in Table 1.

### Cell lines and treatments

Mouse *STHdh*^Q7/Q7^ (CH00097) and *STHdh*^Q7/Q111^ (CH00096) striatal neuronal cells, HD patient-derived fibroblast cell lines [wild type HTT allele/17 CAG repeats, mutant HTT allele/69 CAG repeats (GM04281)] and normal human fibroblast cell line (GM07492) were obtained from Coriell Institute for Medical Research (Camden, New Jersey, USA). Striatal cells were cultured in DMEM high glucose (10566-016, ThermoFisher Scientific), 10% FBS, 5% CO2, at 33°C as described in our previous works[22, 85, 86, 100, 156-158]. Human fibroblasts were maintained at 37°C and 5% CO_2_ in DMEM, high glucose, GlutaMAX supplement supplemented with 10% FBS, 1% penicillin-streptomycin and 1% MEM nonessential amino acids (ThermoFisher Scientific). SUMO1 deletion was carried out in striatal cells using CRISPR/Cas-9 tools obtained from Santa Cruz, as described before[85, 157]. Cycloheximide (CHX, no. C1988, Millipore Sigma) was dissolved in 95% ethanol and used at 75 μM (striatal cells) or 50 μM (human fibroblasts) for indicated time points. To assess autophagy flux, cells were pretreated with chloroquine (CQ, 50 μM, no. C6628, Millipore Sigma) for 4 hr, then proceeded for subcellular fractionation as described below. Stock solution (25 mM) of Ginkgolic acid (GA, no. 6326, Tocris Bioscience) was prepared in DMSO. Striatal cells or human fibroblasts were treated with 100 μM GA and after 12 hrs CHX was added for indicated time points (total time of GA treatment was for 24 hrs)

### Subcellular Fractionation

For subcellular fractionations, mice were euthanized and both striata were dissected immediately and homogenized using a glass dounce homogenizer (5 loose and 5 tight strokes) in buffer A of mitochondria isolation buffer (ThermoFisher Scientific, no. 89874) and kept on ice for 2 minutes. Buffer C was added to each sample and mixed by inverting 5 times. The homogenates were centrifuged at low speed (700g) for 10 minutes to separate nuclei and tissue chunks. The supernatants were immediately loaded on top of 10-50% sucrose gradients and centrifuged at 40000 RPM (SW41Ti rotor) at 4°C for 2 hours. The gradients were fractionated manually (11 or 13 × 1 ml fractions). Using methanol/chloroform, total protein of each fraction was precipitated. The protein pellets were resuspended in 2X LDS buffer and used for western blotting as described below. All the samples were proceeded at the same time. During developing the blots, all the groups were developed at the same time with corresponding antibody. For striatal neuronal cell fractionation, cells were resuspended in buffer A of mitochondria isolation buffer after harvesting and pelleting in PBS. Lysates were kept on ice for 2 minutes and were passed 15 times through a 26-gauge needle and syringe. Lysates were proceeded for subcellular fractionation as mentioned above for striatum samples.

### Western blotting

Striatal tissue were rinsed briefly in PBS and directly lysed in lysis buffer [50 mM Tris-HCl (pH 7.4), 150 mM NaCl, 1% Triton X-100, 1x protease inhibitor cocktail (Roche, Sigma) and 1x phosphatase inhibitor (PhosStop, Roche, Sigma)], sonicated for 2 x 5 sec at 20% amplitude, and cleared by centrifugation for 10 min at 11,000 g at 4°C. Striatal cells or human fibroblasts were lysed in radioimmunoprecipitation assay (RIPA)buffer [50 mM Tris-HCl (pH 7.4), 150 mM NaCl, 1.0% Triton X-100, 0.5% sodium deoxycholate, 0.1% SDS] with 1x complete protease inhibitor cocktail and 1x phosphatase inhibitor, followed by a brief sonication for 2 x 5 sec at 30% amplitude and cleared by centrifugation for 10 min at 11,000g at 4°C. Protein concentration was determined with a bicinchoninic acid (BCA) protein assay reagent (Pierce). Equal amounts of protein (20-50 μg) were loaded and were separated by electrophoresis in 4 to 12% Bis-Tris Gel (ThermoFisher Scientific), transferred to polyvinylidene difluoride membranes, and probed with the indicated primary antibodies. HRP-conjugated secondary antibodies were probed to detect bound primary IgG with a chemiluminescence imager (Alpha Innotech) using enhanced chemiluminescence from WesternBright Quantum (Advansta) The band intensities were quantified with ImageJ software (NIH). Total proteins were then normalized to actin or Gapdh.

### Immunohistochemistry

Immuno-histochemistry was performed as described in our previous work[159, 160]. Briefly, mice were anesthetized and perfused with saline (4°C) and after with 4% PFA (4°C). The whole brain was dissected and postfixed in PFA, cryoprotected by sucrose gradients treatment (10 to 30%) at 4°C for 3 days and embedded in Tissue-Tek. Coronal sections (25 µm thick) were collected on Superfrost/Plus slides and immunostained with the indicated primary antibodies overnight at 4°C. Sections were then incubated with secondary antibodies for 2 h at room temperature (1:250), counterstained with DAPI (Sigma-Aldrich) and mounted with Fluoromount-G mounting medium (ThermoFisher Scientific). Images were acquired by using the Zeiss LSM 880 confocal microscope system with a 20× / 63× objective.

GFAP quantification was made in whole striatal reconstructions from immunostained coronal sections. Using an LSM-800 Zeiss microscope in a tile scan/Z-stack configuration protocol and a 40X objective, we obtained 12 µm thick reconstruction (3 µm stack) from the right and left striatums from each brain section. They consisted of frontal, central, and caudal striatum regions, and about 1 – 3 sections for each striatal region per mouse were observed. GFAP fluorescence intensity was calculated in Z-stack reconstruction for all the groups. Confocal images were analyzed in ImageJ software.

Colocalization coefficient (Mander’s coefficient) analyses and cell quantification for IHC were made in single-plane confocal images to avoid false positive colocalization. Colocalization between DARPP-32 and EM48 was considered intracellular EM48 aggregates, EM48 without DARPP-32 colocalization was deemed extracellular aggregates. EM48 nuclear aggregates were confirmed by DAPI colocalization.

The lateral ventricular area was determined in hematoxylin/eosin-stained sections, from the same animals. Sections at rostral (+1.45 from bregma), central striatum 1 (+1.0 from bregma), central striatum 2 (+0.60 from bregma), and caudal (+0.1 from bregma) lateral ventricles at the striatum level from WT, Q175DN, Q175DN-SUMO1KO and SUMO1KO mice (n = 5 mice per group) were taken using the Leica DM5500B microscope (10x objective). The lateral ventricular area was calculated by analyzing the images using the ImageJ software. N = 5 mice per group, 3 – 4 sections were analyzed and averaged for each striatal region per mouse.

### Immunocytochemistry

Immunostaining was carried out as described in our previous work [100]. Briefly, striatal cells were grown on poly-D-lysine (0.1 mg/ml)–coated glass coverslips and after 24 hours of plating, the medium was changed to Krebs buffer medium devoid of serum and amino acids for 1 hour to induce full starvation conditions. For the stimulation conditions, cells were stimulated with 3mM L-leucine for 15 min. Cells were washed with cold PBS, fixed with 4% paraformaldehyde (PFA; 20 min), treated with 0.1 M glycine, and permeabilized with 0.1% (v/v) Triton X-100 (5 min). After being incubated with blocking buffer [1% normal donkey serum, 1% (w/v) BSA, and 0.1% (v/v) Tween 20 in PBS] for 1 hour at room temperature, cells were stained overnight at 4°C with antibodies against EM48, p62, and LAMP1. Alexa Fluor 488– or Alexa Fluor 568–conjugated secondary antibodies were incubated together with the nuclear stain DAPI for 1 hour at room temperature. Glass coverslips were mounted with Fluoromount-G mounting medium (ThermoFisher Scientific). Images were acquired by using the Zeiss LSM 880 confocal microscope system with a 63× objective.

### Behavioral analysis

#### Open field

An open-field test was used to measure the total activity. Briefly, animals were placed in the center of each square (50 cm × 50 cm) open-top box under bright light and recorded via a ceiling-mounted video camera for 15 min. Locomotor activity was assessed using the EthoVision XT 11.5 animal tracking software (Noldus) and showed the total distance traveled. Before open field performance, all the mice were kept in a waiting room for 30 min. Open-field test was made every two months before Rotarod and beam walk evaluation.

#### Rotarod

The accelerating rotarod test was used to quantify motor alterations in all the genotypes. Before evaluation day, all the mice were trained for 3 min in a no-accelerating rod (20 rpm) for two consecutive days. On the day of the test, all the mice were challenged in an accelerating rod (4 to 40 rpm, with a cut off of 5 min), and the average of the latency to fall for three trials was used for the analysis (20 min of resting were used between each trial). Rotarod was evaluated every two months old, starting at two months after born.

#### Beam walk

Alterations of the motor coordination also were evaluated using the beam walk test. The beam walk system consisted of the suspended beam (100 cm long) at 50 cm high with a chamber (12×12×12 cm) at the end of a 100 cm long beam. Every two months, all the mice were trained for three days to cross a beam. On the first day of the training, we used a 12 mm width beam; for the second day, we used a 9 mm width beam, and on the third day (evaluation day), a 6 mm width beam was used. For each evaluation session, three trials were made with 10 min between them. The average time to cross the beam (100 cm) was recorded on the evaluation day. Before the trials and between them, the mice were kept in the beam walk chamber.

#### Behavioral battery evaluation

The behavior battery evaluation was done at 13 months old; each test was performed in triplicate, measured ledge walking, clasping, gait, kyphosis, and tremor. Individual measures were scored on a scale of 0 (absence of relevant phenotype) to 3 (most severe manifestation), as described previously [100, 161], with the addition of testing for tremor. To determine tremor, mice were placed in a clean cage and observed for 30 seconds. Each mouse was scored as follows: 0, no signs of tremor; 1, present but mild tremor; 2, severe intervals of tremor or constant moderate tremor; 3, extreme chronic tremor. The composite score was generated as the mean of all five behavioral battery tests.

### RNA seq

Three striatum samples from each genotype (WT, Q175DN, Q175DN-SUMO1KO and SUMO1KO) were lysed in Trizol reagent. RNA was extracted from miRNeasy kit from Qiagen (cat. No. 217004) and RNA-seq and IPA were performed in Scripps Genomics and Bioinformatics Core, Jupiter, FL, USA as described[22].

### Statistical analysis

Data are presented as mean ± SEM as indicated. Except where stated all experiments were performed at least in biological triplicate and repeated at least twice. The mouse behavioral and the data analysis was carried out in a blinded manner. Images were quantified using ImageJ (FIJI). Behavioral data consisted of categorical and continuous outcomes. Categorical data was analyzed using Fisher’s Exact test. Statistical comparison was performed between groups using two-tailed Student’s t-test, one-way analysis of variance (ANOVA) followed by Tukey’s multiple comparison test or Bonferroni’s multiple comparisons test and two-way ANOVA or repeated measure two-way ANOVA or two-way ANOVA mixed-effects model (REML) followed by Tukey’s multiple comparison test or Bonferroni post-hoc test as indicated in the figure legends. Significance was set at p < 0.05. All statistical tests were performed using Prism 9.0 (GraphPad software).

## Data availability

Sequencing data have been submitted to the Gene Expression Omnibus (GEO) data repository.

## Acknowledgments

We would like to thank Melissa Benilous for her administrative help and the members of the lab for their continuous support and collaborative atmosphere. We like to thank members at the Scripps proteomics and genomic core for their help and expertise. This research was partially supported by a training grant in Alzheimer’s drug discovery from the Lottie French Lewis Fund of the Community Foundation for Palm Beach and Martin Counties; grant awards from NIH/NINDS R01-NS087019-01A1, NIH/NINDS R01-NS094577-01A1, a grant from Cure for Huntington Disease Research Initiative (CHDI) foundation and Scripps bridge funding.

## Author contributions

S.S and N.S conceptualized the project and co-designed experiments with U.N.R.J. who performed mouse behavior experiments, immunohistochemistry, immunostaining, confocal, colocalization analysis and analyzed the data. M.S., U.N.R.J. and N.S carried out SUMO deletion and Western blotting experiments and quantification. S.S and N.S and wrote the manuscript with inputs from coauthors.

## FIGURE LEGENDS

**Supplementary Figure 1.**
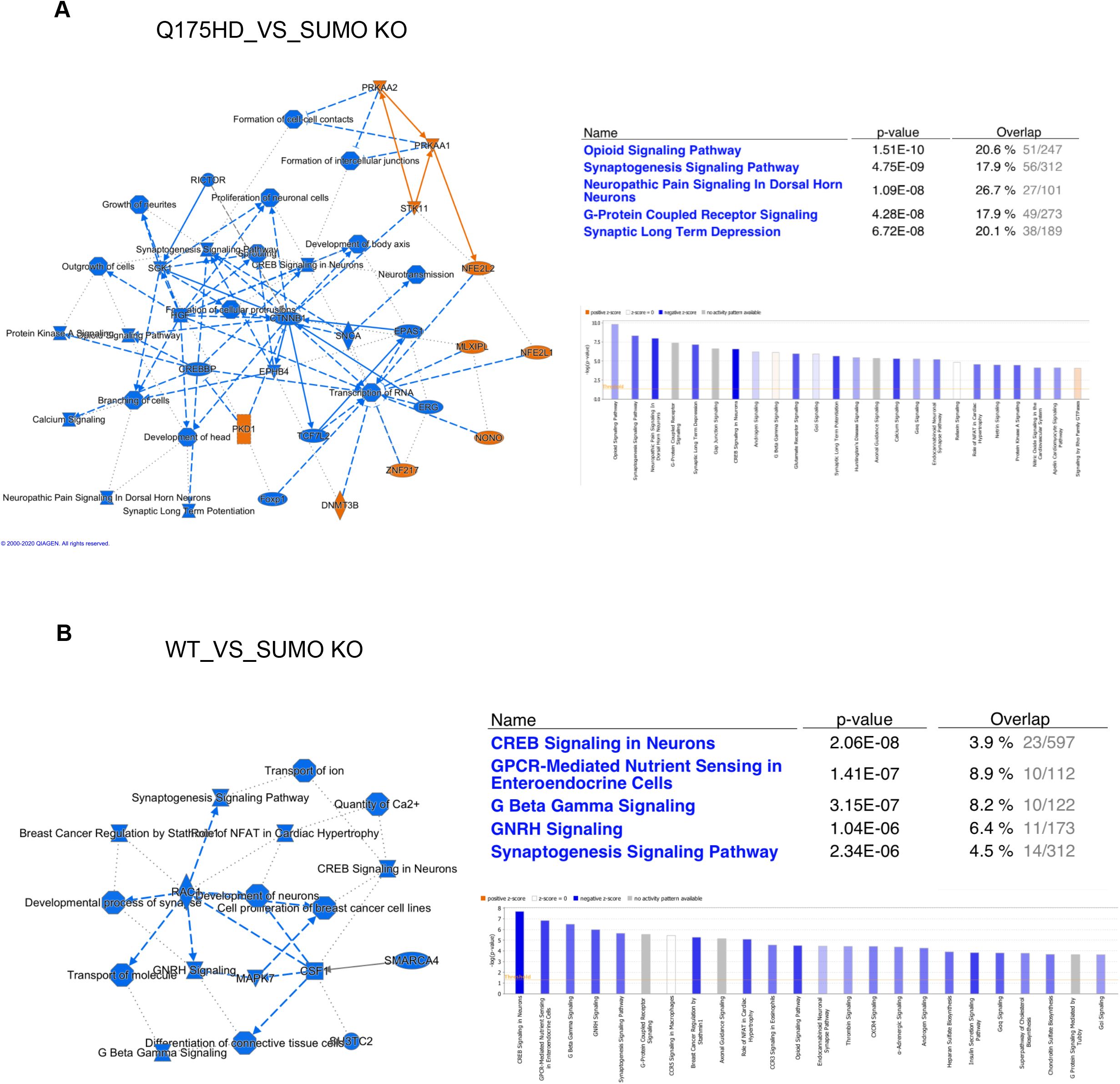
(A, B) IPA network analysis between WT and SUMO1KO mice and SUMO1KO and Q175DN mice.

